# NETGE-PLUS: standard and network-based gene enrichment analysis in human and model organisms

**DOI:** 10.1101/750661

**Authors:** Samuele Bovo, Pier Luigi Martelli, Pietro Di Lena, Rita Casadio

**Author notes:** **Corresponding Author** (PLM).

## Abstract

Omics techniques provide a spectrum of information that needs to be disentangled to characterize complex traits at the molecular level. The gap between genotype and phenotype must be closed by reconciling the genome information with the set of molecular pathways and biological processes describing the phenotype. In dealing with this problem, gene enrichment analysis has become the most widely adopted strategy. Here, we present NETGE-PLUS, a web-server for standard and network-based functional interpretation of gene sets of human and of model organisms, including *S. scrofa, S. cerevisiae, E. coli* and *A. thaliana*. NETGE-PLUS enables the functional enrichment of both simple and ranked lists of genes, also introducing the possibility of exploring relationships among KEGG pathways. A web interface makes data retrieval complete and user-friendly. NETGE-PLUS is publicly available at http://net-ge2.biocomp.unibo.it

## INTRODUCTION

The analysis of large-scale genomic and proteomic data, aimed at characterizing complex traits at the molecular level, routinely returns lists of “relevant” genes/proteins. Over-representation analysis (ORA) is a widely adopted method to endow a gene set with functional annotations, with the aim of disentangle the complex relationships among genes, their functions, and the phenotype of interest. ORA tests whether some biological features are significantly more frequent in the given gene set than expected by chance. Alternatively, when genes are endowed with some numerical score (e.g. the fold change of differential expression), Gene Set Enrichment Analysis (GSEA) allows including the weighted contribution of a large ranked list of genes to perform functional characterization [1].

Previously we described NET-GE, a tool for performing ORA on sets of human genes, considering annotations associated to each gene and functional features, derived from the analysis of the human interactome [2–3]). Exploiting the information contained in biological networks [4], NET-GE enriches Gene Ontology (GO) terms [5], KEGG PATHWAY [6] and Reactome [7] annotations. However, NET-GE does not allow GSEA, and it is limited to human genes and their related interactions. NET-GE adopts the KEGG BRITE hierarchy (http://www.genome.jp/kegg/kegg3b.html) that provides a logical organization more than a complete description of the proximity relationships among pathways.

Despite cross species comparisons have shown conserved patterns of protein interactions [8], interactomes remain species specific.

Here we introduce NETGE-PLUS, a new web-server that allows computing gene enrichment analysis not only in humans, but also in organisms widely adopted in translational, biomedical and biotechnological research, including *S. scrofa, E. coli, S. cerevisiae* and *A. thaliana*. NETGE-PLUS allows to perform gene set enrichment analysis also when genes are ranked according to some criteria (e.g. expression level), implementing both the ORA and GSEA procedures. Furthermore, by introducing links among different KEGG maps, it is possible to define a network of pathways (KEGG-NET) for an approximate description of the possible complexity underlying the set of given genes.

## MATERIAL AND METHODS

### Databases

NETGE-PLUS includes the interactomes of *H. sapiens* (hsa; taxid:9606), *S. scrofa* (ssc; taxid:9823), *S. cerevisiae* (sce; taxid:4932), *E. coli* (eco; taxid:51145) and *A. thaliana* (ath; taxid:3702), as derived from STRING v.10.5 [9]. The identifiers adopted in the procedure are: (i) the ENSEMBL protein identifiers, for *H. sapiens* and *S. scrofa* and (ii) the systematic locus identifiers, adopted for *S. cerevisiae, E. coli* and *A. thaliana*. All the links with a combined STRING score ≥ 0.4 (medium confidence) were retained irrespectively of the supporting evidence. The modules have been computed for the Gene Ontology terms (v.169; https://www.ebi.ac.uk/GOA), KEGG (v.83.2; http://www.kegg.jp/) and Reactome (v.61; https://reactome.org/) pathways.

### Processing the interactomes

NETGE-PLUS relies on modules of functionally related genes, pre-computed as described by Di Lena et al. [2]. Briefly, each module is built starting from (i) a set of genes sharing a specific functional term (seed set) and (ii) an interactome. Each seed set is extended into a compact and connected module of interacting proteins by computing all the shortest paths among the seeds genes. By applying measures based on graph and information theory, modules are reduced into minimal connecting networks while preserving the distances among seeds. The resulting modules, containing seed nodes and some of their interacting partners (connecting nodes), are at the basis of the functional enrichment. Details about the module construction can be found in the Supplementary Informations and online (see the help web-page).

### Enrichment procedures

NETGE-PLUS implements both a standard and a network-based gene enrichment analysis. Given an input gene/protein set, genes are mapped into the subnetworks of each annotation database. NETGE-PLUS allows to perform: (i) ORA, thorough the Fisher’s exact test and (ii) GSEA, through a Kolmogorov–Smirnov-like statistic, as described by Subramanian et al. [1] and re-implemented using the Python2.7 package GSEAPY (v0.9.4; https://pypi.python.org/pypi/gseapy). For GSEA we use a weighted enrichment scoring statistic and 100 permutations. Standard enrichment analysis includes only annotations of seed nodes in each subnetwork while the network-based one includes seeds and their connecting nodes. For multiple testing correction the user can select either the Bonferroni or the Benjamini-Hochberg (False Discovery Rate, FDR) procedures [10].

### Implementation of the web server

The web server runs on a web2py engine (http://www.web2py.com/) and it is optimized to work with all common web browsers. The analysis runs asynchronously: upon request submission, the server displays a book-markable page that is periodically updated until the job completion. A link to the result page is given to the user as soon as the job is completed.

The final visualization of the results exploits the Graphviz library (http://www.graphviz.org/) for laying out the acyclic directed graphs for both Gene Ontology, KEGG and REACTOME. KEGG-NET results are rendered via the JavaScript library Cytoscape.js (http://js.cytoscape.org/). Enriched terms from these annotation systems are highlighted. In addition, the Web server shows dynamic network renderings, based on the JavaScript library d3.js (http://d3js.org/), for the visualization of the underlying interaction networks involving a specific term and the network of pathway.

For multiple submissions, each request is queued, and it runs as soon as there is available computing power. Running time depends on size of the input set (from two up to 200; higher numbers may require a long time) and ranges from 1 to 10 minutes.

The user can also provide an e-mail address used to e-mail him/her as soon as results are ready.

### Study cases

Three gene sets, related to human and porcine diseases, were investigated to qualitative evaluate the performance of NETGE-PLUS. Here, we focus only on one of them, the Non-alcoholic Fatty Liver Disease (NAFLD) study case. The other two study cases are presented in the Supplementary Information and online (see the tutorial web-page).

We retrieved from Phenopedia [11] a list of genes that possibly contribute to the development of NAFLD. Among the 408 NAFLD-related genes, we selected the ones supported at least from five publications. As result, we obtained a list of 28 genes. This gene set was investigated by NETGE-PLUS by performing ORA over the KEGG-NET resource. We considered statistically enriched terms with a *p*-value < 0.01, after the correction with the Bonferroni procedure.

## RESULTS AND DISCUSSION

### NETGE-PLUS web server

NETGE-PLUS accepts different gene names and identifiers (UniProtKB, ENSEMBL gene and protein, official gene name and the systematic locus identifier), plus the related scores, in the case of GSEA. As output, NETGE-PLUS returns two tables (one for the standard method and one for the network-based method; Figure 1A) listing the over-represented terms. Tables report basic information (e.g. term name, *p*-value(s), input genes) plus other statistics that allow a better understanding of the specificity of enriched terms. These include the information content (IC), a measure of function specificity [12], and the level in the ontology hierarchy. The results page provide also graphs depicting (i) the functional connection among enriched terms (the hierarchy or the network of pathways; Figure 1B) and (ii) the organization of functional modules (sub-networks; Figure 1C). Tables and graphs can be downloaded and locally managed.

**Figure 1.**
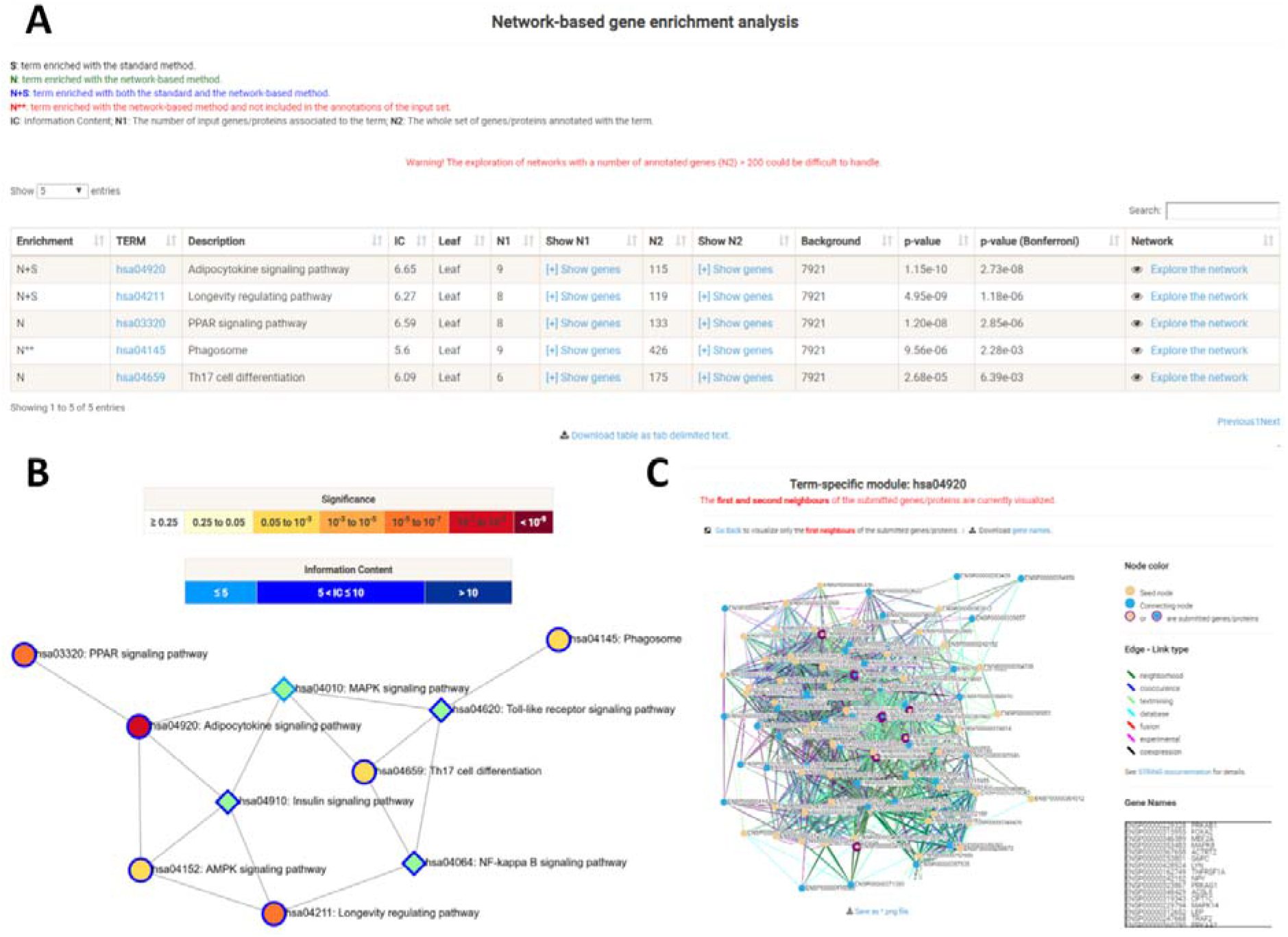
NETGE-PLUS result pages for the NAFLD case study. (A) Table of network-based over-represented KEGG-NET pathways. (B) Network of pathways. Circles represent enriched pathways while diamonds the connecting pathways. Circle colour represents the magnitude of enrichment, while the green colour of diamonds highlights the presence of at least one input genes associated to them. (C) Functional module of the over-represented pathway hsa04920. Seed nodes (yellow) represents genes directly annotated with the term while connecting nodes (blue) represents connecting genes. Nodes presenting a purple border identifies the part of the submitted genes. The link type is highlighted based on the seven different channels of STRING.

Input and output details are presented in the Supplementary Information. Additional details are also available online (see the help web-page).

### Understanding the functional relationships among over-represented KEGG pathways

When enriching functional pathways, it is useful to represent them in a larger context that comprises the most related pathways. While for Reactome a full hierarchy is provided, in the case of KEGG, the BRITE hierarchy gives only a categorization of the different KEGG maps. In order to better understand the overall organization of all the pathways, we exploit the information contained in the links among different KEGG maps by defining KEGG-NET. When a set of genes enriches different KEGG pathways, the most related connecting pathways are determined by computing the pairwise shortest paths through the links. Since the pathway network is highly connected, we retain only the paths with a maximum length equal to 2 (no. of edges). KEGG map01100 (whole metabolism) and the disease-related maps are not considered in this procedure, because of their dense connectivity. In this way, NETGE-PLUS users have the possibility to quickly understand which biological dependences exist among the enriched pathways.

### Non-alcoholic fatty liver disease: a case study involving the KEGG-NET resource

In the following, we deal with a specific case study that is also described in the online tutorial. We make use of the KEGG-NET resource to dissect the biological complexity of the Non-alcoholic Fatty Liver Disease (NAFLD). Defined as a genetic-environmental-metabolic stress-related disease, NAFLD is a pathology characterized by an excessive fat accumulation in the liver even in the absence of alcohol consumption. NAFLD encompasses a spectrum of diseases, from simple steatosis to non-alcoholic steatohepatitis (NASH), which can progress to cirrhosis and hepatocellular carcinoma. There are also increasing evidences that NAFLD represents the hepatic component of a metabolic syndrome characterized by obesity, hyperinsulinemia, peripheral insulin resistance, diabetes, hypertriglyceridemia and hypertension [13]. Moreover, several studies have identified many genetic variations that may be associated with the development of NAFLD [14].

The analysis of the 28 NAFLD-related genes highlighted a total of six over-represented pathways (Table 2), three detected via the standard enrichment analysis while other three added by the network-based procedure.

**Table 1.**
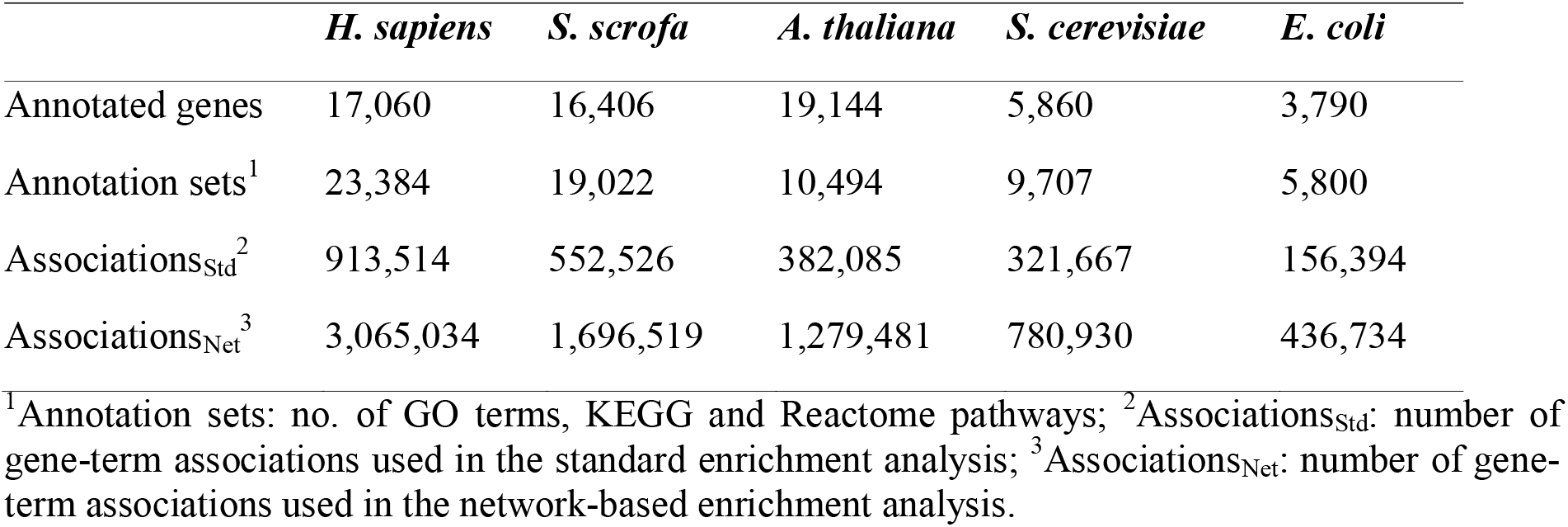
NETGE-PLUS summary statistics.

**Table 2.** Non-alcoholic fatty liver disease case study. Gene enrichment analysis over the KEGG-NET resource.

| Enrichment<sup>1</sup> | Term<sup>2</sup> | N1<sup>3</sup> | N2<sup>4</sup> | Background<sup>5</sup> | Bonferroni<sup>6</sup> | Description<sup>7</sup> |
| --- | --- | --- | --- | --- | --- | --- |
| S | hsa04920 | 6 | 69 | 6,516 | 8.44E-05 | Adipocytokine signaling pathway |
| S | hsa04211 | 6 | 90 | 6,516 | 4.12E-04 | Longevity regulating pathway |
| S | hsa04152 | 6 | 121 | 6,516 | 2.32E-03 | AMPK signaling pathway |
| N | hsa03320 | 8 | 133 | 7,921 | 2.85E-06 | PPAR signaling pathway |
| N** | hsa04145 | 9 | 426 | 7,921 | 2.28E-03 | Phagosome |
| N | hsa04659 | 6 | 175 | 7,921 | 6.39E-03 | Th17 cell differentiation |

<sup>1</sup>Enrichment: Standard (S) and Network-based (N) procedure. N\*\* indicates a new enriched term not directly associated to the input gene/proteins; <sup>2</sup>Term: functional annotation identifier; <sup>3</sup>N1: Input genes/proteins belonging to the term; <sup>4</sup>N2: genes associated to the functional term; <sup>5</sup>Background: number of genes used as background at the Fisher's exact test; <sup>6</sup>Bonferroni: p-value corrected by using the Bonferroni procedure; <sup>7</sup>Description: brief explanation of the term.

The pathways over-represented by the standard method are: the Adipocytokine signaling pathway, the longevity regulating pathway and the AMPK signaling pathway. Considering the graph generated by linking the whole set of enriched pathways (Figure 1B), the three pathways are connected within a chain. All of them are also linked to the insulin signalling pathway (not included in the enriched set). Insulin resistance plays an important role in NAFLD and it is caused by adipocytokines, a specific kind of cytokines secreted by the adipose tissue. Among them, adiponectin is an anti-inflammatory and anti-diabetic adipocytokine that exerts its actions by the activation of adenosine monophosphate (AMP)-activated kinase (AMPK) and PPARα [15]. Interestingly, the PPAR signalling pathway is one of the terms enriched with the network based procedure.

Another important cytokine within the adipocytokine pathway is leptin. It binds the leptin receptor (LEP-R) and triggers a phosphorylation chain resulting in the activation of the MAPK pathway [16]. This is another connecting pathway in the network. One of the members of the MAPK pathway, namely the protein kinase c-Jun N-terminal kinase (JNK), is closely related to insulin resistance. Moreover, rat models with activated JNK present phenotypes related to NAFLD, such as hepatocyte fat accumulation and cell injury [13].

In the graph presented in Figure 1B, the MAPK pathway links the adipocytokine and insulin signalling pathways with the Th17 cell differentiation pathway (enriched with the network-based procedure). Interestingly, Th17 cells have been associated with hepatocellular steatosis and inflammatory processes via the production of IL-17, that is also implicated in insulin resistance.

Moreover, secretion of IL-17 is triggered and perpetuated through the nuclear factor-κB (NF-κB) [17], and the NF-κB pathway is a connecting node.

The last term enriched with the network procedure is “Phagosome”, that is linked to the other nodes through the “Toll-like receptor signalling pathway”. Both these terms computationally derived with NETGE-PLUS suggest the involvement of macrophages in NAFLD, in particular in relation to the reprogramming induced by cytokines. The role of macrophages in NAFDL from initial steatosis to advanced fibrosis has been previously reviewed in Krenkel and Tacke [18]

In conclusion, by highlighting the links among the different metabolisms, KEGG-NET clearly helps the user to globally understand the biology at the basis of the phenotype of interest.

## CONCLUSIONS

In this article we presents NETGE-PLUS web server, the new version of our network-based gene enrichment analysis tool. NETGE-PLUS arrives with major improvements that include: (i) updated annotations for H. sapiens, (ii) the possibility to operate with five well-studied model organisms (*S. scrofa, S. cerevisiae, E. coli and A. thaliana*), (iii) the possibility to perform GSEA, (iv) the possibility to understand the functional relationships among over-represented KEGG pathways via the KEGG-NET resource and (v) a restyled web interface.

As result, within a user friendly environment, NETGE-PLUS makes network-based gene enrichment analysis (ORA and GSEA) accessible to researchers working with the main model organisms encompassing the different kingdoms of life. Moreover, through the analysis of the links established by biologically relevant pathways (via the KEGG-NET resource), NETGE-PLUS allows to disentangle biological complexity.

## ASSOCIATED CONTENT

### Supplementary Information

workflow of the gene enrichment analysis, web-server details and case studies.

## Author Contributions

SB implemented the web application, performed enrichment analyses and drafted the manuscript. PLM and PDL supervised the module generation. RC and PLM supervised the project and drafted the manuscript. All authors read and approved the final manuscript.

## Notes

The authors declare no competing financial interest. NETGE-PLUS is publicly available at http://net-ge2.biocomp.unibo.it

---

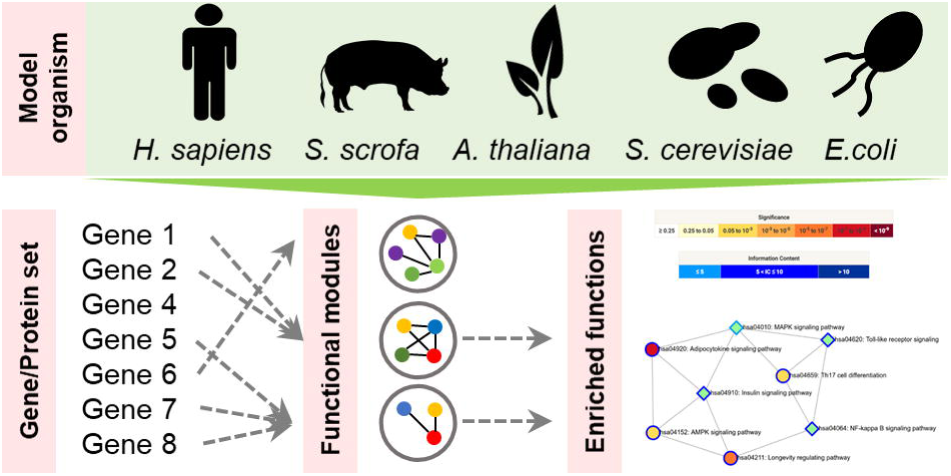

