## Supplementary Information for "NETGE-PLUS: standard and network-based gene enrichment analysis in human and model organisms"

### 1. Workflow of the gene (set) enrichment analysis.

Given a protein/gene set, NETGE-PLUS implements both a standard and a network-based gene enrichment analysis. Both procedures can be adopted when running either a classic over-representation analysis (ORA; via a Fisher's exact test) or a gene set enrichment analysis (GSEA [1]; via a Kolmogorov–Smirnov-like statistic). Gene enrichment analyses can be performed, separately, over the three branches of the Gene Ontology resource [2] (http://www.geneontology.org/; Biological process – GO-BP, Molecular Function – GO-MF, Cellular Component – GO-CC), the KEGG PATHWAY [3] (http://www.kegg.jp/) and Reactome [4] (https://reactome.org/) databases. Moreover, NETGE-PLUS introduces also KEGG-NET, an in-house derived version of KEGG PATHWAY that neglects the KEGG BRITE hierarchy (http://www.genome.jp/kegg/kegg3b.html) and introduces links among pathways on the basis of shared proteins and metabolites (map links, as provided by the KEGG maps themselves).

Statistics on the number of annotated genes (background) and annotating features, for each database and organism, are provided online (http://net-ge2.biocomp.unibo.it/enrich/default/statistics).

The network-based procedure relies on modules of functionally related genes, precomputed as described in Di Lena *et al.* [5]. Briefly, each module is built starting from (i) a set of genes related to a specific functional term (seed set) and (ii) the STRING v.10.5 interactome [6] (https://version-10-5.string-db.org/). Each seed set is extended into a compact and connected module of the interacting proteins by computing the whole set of shortest paths among the seed genes. By applying measures based on graph and information theory, modules are then reduced into minimal connecting networks while preserving the distances among seeds. The resulting modules, containing seed nodes and some of their interacting partners (connecting nodes), are at the basis of the network-based functional enrichment. An outline of the module construction is given in Supplementary Fig. S1.

It is important to note that the standard procedure can enrich only functional terms already associated to the input genes/proteins. On the contrary, the network-based procedure allows to detect statistical associations with functional terms not included in the annotations of the input set. These "new" terms represent one of the main added values of the network-based enrichment analysis.

Moreover, the GSEA procedure allows to weight the individual contribute of each input gene, by ranking it with a relevance value provided by the user. For example, by providing for each gene the corresponding fold change as derived from a differential expression experiment, it is possible to extend the analysis to all genes of an organism, without the need of filtering them in advance.

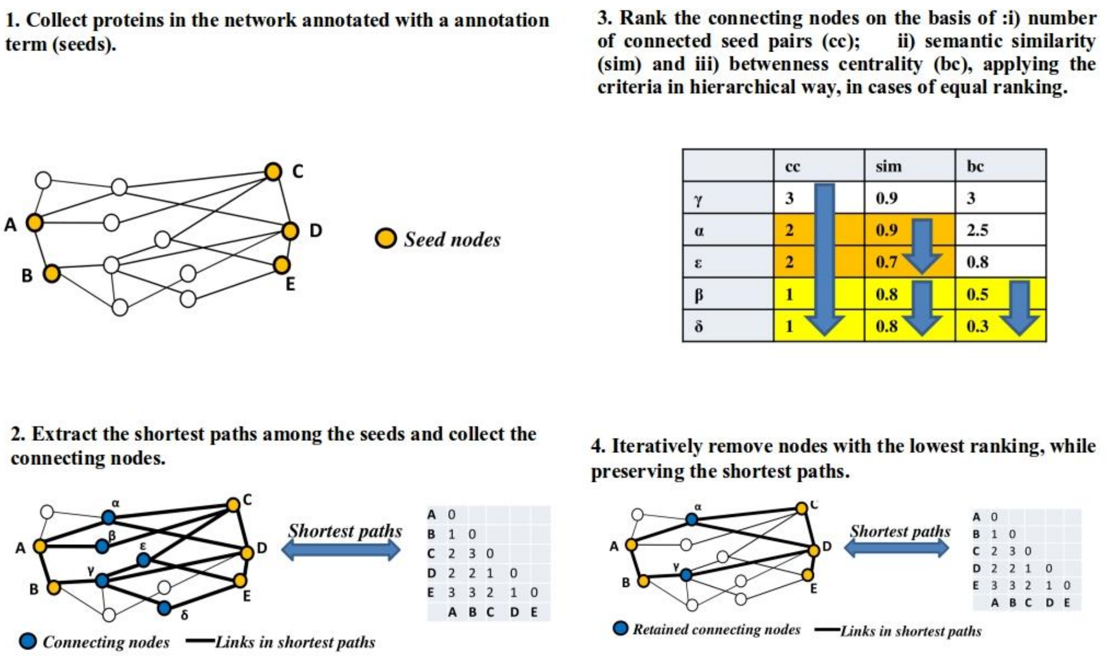


**Supplementary Figure S1.** Outline of the module generation in NETGE-PLUS. Figure adapted from Di Lena *et al. [5]*.

### 1.1 NETGE-PLUS input

NETGE-PLUS accepts UniProtKB accessions (UNIPROT ACC), ENSEMBL IDs (genes and proteins), gene names and the systematic locus ID. Examples of identifiers for the different organisms are provided in Supplementary Table S1. However, the usage of gene names is discouraged because of the ambiguous association with identifiers. Moreover, in the GSEA mode, it is necessary to provide a numerical relevance score for each input gene/protein.

The web interface allows the user to select: 1) the organism of interest, 2) the gene/protein identifier, 3) the annotation database (GO terms, KEGG and Reactome pathways), 4) the analysis method (ORA or GSEA), 5) the multiple testing correction procedure (Bonferroni or Benjamini-Hochberg; [7]) and 6) the significance threshold. An e-mail address can be provided in order to be notified as soon as results are ready.

**Supplementary Table S1.** Examples of identifiers for the asparagine synthase (glutamine-hydrolysing) [EC:6.3.5.4].

| Identifier | *H. sapiens* | *S. scrofa* | *S. cerevisiae* | *E. coli* | *A. thaliana* |
|---|---|---|---|---|---|
| **UNIPROT_ACC** | P08243 | D0G0C6 | P49089 | P22106 | P49078 |
| **ENSEMBL_GENE_ID** | ENSG00000070669 | ENSSSCG00000015340 | YPR145W | b0674 | AT3G47340 |
| **ENSEMBL_PROTEIN_ID** | ENSP00000175506 | ENSSSCP00000016267 | YPR145W | b0674 | AT3G47340.1 |
| **STANDARD_GENE_NAME^** | ASNS | ASNS | ASN1 | asnB | ASN1 |
| **SYSTEMATIC_GENE_NAME*** | - | - | YPR145W | b0674 | AT3G47340 |

^ The gene nomenclature follows the standards for the different organisms; * The SYSTEMATIC_GENE_NAME has been introduced as identifier for *E. coli*, *S. cerevisiae* and *A. thaliana*.

### 1.2 NETGE-PLUS output

NETGE-PLUS gives as output two tables listing the enriched terms: one table for the standard method and one table for the network-based method. Each table reports: (i) the identifier of the functional term (externally linked to the source database), (ii) the description of the term, (iii) the information content (IC) [8] of the term, (iv) the level in the ontology hierarchy (leaf of ancestor), (v) the number and the list of submitted genes/proteins associated to the term, (vi) the number and the list of genes/proteins annotated with the term in the whole organism-specific dataset, (vii) the number of genes used as background and (vii) the raw and corrected *p*-values. Genes are linked to the corresponding Uniprot entry.

An extra column reporting a link for the module visualization web-page is available for the network-based enrichment. NETGE-PLUS allows the inspection of each functional module. Modules are presented in a STRING-like manner by highlighting the different link types (STRING channels; see https://string-db.org/help/getting_started/#evidence). Genes directly associated to the functional term, connecting genes and input genes are presented in different colours. Nodes and arcs, with the related information, can be downloaded for an in-house inspection. When performing GSEA, a link to the enrichment plot is also provided.

### 2. Case studies

In the following, we briefly present two case studies demonstrating how NETGE-PLUS can help in the interpretation and understanding of gene sets related to diseases.

The first case study – approached with a classic ORA procedure – deals with a gene set related to a human disease as retrieved from the OMIM resource (https://www.omim.org/).

The second case study – approached with the GSEA procedure – considers a set of genes found differently expressed in a pig cohort exposed to a viral infection.

### 2.1 ICHTHYOSIS, CYCLIC, WITH EPIDERMOLYTIC HYPERKERATOSIS (CIEHK)

Cyclic ichthyosis with epidermolytic hyperkeratosis (CIEHK) is a subtype of bullous congenital ichthyosiform erythroderma (BCIE). Based on its clinic-pathological feature – annular and polycyclic erythematous plaques over the proximal extremities and trunk – the disease has been named also as “annular epidermolytic ichthyosis” [9]. CIEHK involves cornification caused by mutations in the keratin 1 gene (KRT1; HGNC:6412; ENSG00000167768) or keratin 10 (KRT10; HGNC:6413, ENSG00000186395) gene.

We performed standard and network-based ORA on the two genes and we retained enriched terms with a *p*-value < 0.01, after correction with the Benjamini-Hochberg procedure. For the sake of clarity, by considering the hierarchical structure of the annotation sources, we report in Supplementary Tables S2, S3 and S4 only over-represented leaf terms.

Over the GO-BP resource (Supplementary Table S2), the standard method highlights processes strictly related to the disease: “peptide cross-linking”, “skin epidermis development”, “cornification” and “keratinization”. The network-based method adds three new terms: “skin development”, “regulation of keratinocyte proliferation” and “neuromuscular process controlling posture”. Interestingly, “skin development” is strictly related to the “cornification” process, and “keratinocyte proliferation” has been observed in mouse model for BCIE [10]. Moreover, looking for the involvement of neuromuscular processes, a psycho-motor retardation (motor skills and language) has been observed in a subject suffering of ichthyosis (because of a mutation in the KRT10 gene; [11]).

**Supplementary Table S2.** CIEHK case study: over-represented GO-BP.

| Enrichment[1] | Term[2] | N1[3] | N2[4] | Background[5] | FDR[6] | Description[7] |
|---|---|---|---|---|---|---|
| S | GO:0098773 | 2 | 10 | 15699 | 2,37E-05 | skin epidermis development |
| S | GO:0018149 | 2 | 59 | 15699 | 4,51E-04 | peptide cross-linking |
| S | GO:0070268 | 2 | 113 | 15699 | 1,11E-03 | cornification |
| S | GO:0031424 | 2 | 199 | 15699 | 2,08E-03 | keratinization |
| N** | GO:0050884 | 2 | 46 | 17237 | 1,01E-03 | neuromuscular process controlling posture |
| N** | GO:0010837 | 2 | 109 | 17237 | 3,83E-03 | regulation of keratinocyte proliferation |
| N** | GO:0043588 | 2 | 173 | 17237 | 5,81E-03 | skin development |

[1]Enrichment: Standard (S) and Network-based (N) procedure. N** indicates a new enriched term not directly associated to the input gene/proteins; [2]Term: functional annotation identifier: [3]N1: Input genes/proteins belonging to the term; [4]N2: genes associated to the functional term; [5]Background: number of genes used as background at the Fisher's exact test; [6]FDR: *p*-value corrected by using the Benjamini-Hochberg (False Discovery Rate, FDR) procedure; [7]Description: brief explanation of the term.

Over the GO-CC resource (Supplementary Table S3), the standard method enriches the term "cornified envelope". The network-based method adds "keratin filament", and the new term "desmosomes". Their involvement seems coherent with the features of the pathology, since desmosomes participate in the formation of the "cornified envelope" [12].

**Supplementary Table S3.** CIEHK case study: over-represented GO-CC.

| Enrichment | Term | N1 | N2 | Background | FDR | Description |
|---|---|---|---|---|---|---|
| S | GO:0001533 | 2 | 63 | 12043 | 3.23E-04 | cornified envelope |
| N** | GO:0030057 | 2 | 56 | 15476 | 6.17E-04 | desmosome |
| N | GO:0045095 | 2 | 276 | 15476 | 5.07E-03 | keratin filament |

Columns descriptors are given as in **Supplementary Table S2**. N** indicates a new enriched term not directly associated to the input gene/proteins.

Over the Reactome database, the standard method highlights the pathway "Formation of the cornified envelope" (Supplementary Table S4). As previously reported, its over-representation seems coherent with the features of the disease. Moreover, the network-based enrichment analysis (Supplementary Table S4) adds the new term "Post-translational modification: synthesis of GPI-anchored proteins". Interestingly, epidermal-specific impairments of GPI anchor in mice defective of Pig-a gene (essential for the formation of the GPI anchor) include harlequin ichthyosis-like features [13].

**Supplementary Table S4**. CIEHK case study: over-represented Reactome pathways.

| Enrichment | Term | N1 | N2 | Background | FDR | Description |
|---|---|---|---|---|---|---|
| S | R-HSA-6809371 | 2 | 129 | 10248 | 7.86E-004 | Formation of the cornified envelope |
| N** | R-HSA-163125 | 2 | 374 | 13399 | 5.44E-003 | Post-translational modification: synthesis of GPI-anchored proteins |

Columns descriptors are given as in **Supplementary Table S2**. N** indicates a new enriched term not directly associated to the input gene/proteins.

### 2.2 Effect of the Pseudorabies Virus infection on the swine transcriptome.

Pseudorabies virus (PRV) is a swine neurotropic virus that causes: (i) fetal encephalitis in new-born pig, (ii) respiratory disorder in fattening pigs and (iii) reproductive failure in sow. To characterize the host-virus interactions, Miller *et al.* [14] investigated the effect of a PRV infection on the transcriptome of tracheobronchial lymph nodes (TBLN) over the time (at 1, 3, 6 and 14 days). The authors performed a Digital Gene Expression Tag Profiling of RNA isolated from draining TBLN from a total of 40 pigs, either clinically infected with a PRV (no. 20 pigs) or uninfected (no. 20 pigs). By comparing the gene expression profiles, differentially expressed (DE) genes were detected. Biological processes and pathways involving the sets of DE genes were investigated by the authors by adopting the GSEA procedure.

By using GSEA, Miller *et al.* [14] observed the induction of the host innate immune response, including the upregulation of interferon responsive genes, inflammatory response genes, and cytokine–cytokine receptor interaction.

Considering the DE genes at time 1, we applied the GSEA-based procedure implemented in NETGE-PLUS, taking advantage of the network-derived modules. The analysis was carried out considering statistically enriched terms with a *p*-value < 0.05, after the correction with the Benjamini-Hochberg procedure. This is much lower than the 0.25 threshold often adopted in GSEA analysis. For the sake of clarity, by considering the hierarchical structure of the annotation sources, we report in Supplementary Table S5 only over-represented leaf terms.

Over the GO-BP resource, out of the 143 submitted genes, 133 were present in NETGE-PLUS. A total of 110 and 122 genes were effectively included in the seed sets and in the network-based modules, respectively. The standard method highlights the following processes (Supplementary Table S5): “response to virus” and “carboxylic acid biosynthetic process”. The use of the network lead to add 9 terms strictly related to the immune system and correctly highlighting the immunogenic effect of the virus (Supplementary Table S5). Thus, NETGE-PLUS directly pointed out the activation of immunological processes involved in the response to virus, and like Miller *et al.* [14] gene sets related to the cytokines and other molecular mediators of immune response were here evidenced.

**Supplementary Table S5.** PRV case study. GSEA over the GO-PB.

| Enrichment | Term | N1 | N2 | Background | FDR | Description |
|---|---|---|---|---|---|---|
| S | GO:0046394 | 6 | 145 | 13024 | 1.08E-02 | carboxylic acid biosynthetic process |
| S | GO:0009615 | 9 | 123 | 13024 | 1.88E-02 | response to virus |
| N | GO:0030098 | 10 | 348 | 15571 | 3.89E-02 | lymphocyte differentiation |
| N | GO:0002702 | 5 | 114 | 15571 | 4.00E-02 | positive regulation of production of molecular mediator of immune response |
| N | GO:0002824 | 8 | 138 | 15571 | 4.03E-02 | positive regulation of adaptive immune response based on somatic recombination of immune receptors built from immunoglobulin superfamily domains |
| N | GO:0045089 | 9 | 234 | 15571 | 4.13E-02 | positive regulation of innate immune response |
| N | GO:0051224 | 11 | 410 | 15571 | 4.19E-02 | negative regulation of protein transport |
| N | GO:0051048 | 8 | 320 | 15571 | 4.54E-02 | negative regulation of secretion |
| N | GO:0045087 | 17 | 684 | 15571 | 4.72E-02 | innate immune response |
| N | GO:0002718 | 3 | 90 | 15571 | 4.75E-02 | regulation of cytokine production involved in immune response |
| N | GO:0002703 | 5 | 324 | 15571 | 4.75E-02 | regulation of leukocyte mediated immunity |
| N+S | GO:0051607 | 16 | 247 | 15571 | 4.96E-02 | defense response to virus |

Columns descriptors are given as in **Supplementary Table S2**.
